## Supplementary Info for "Gapless provides combined scaffolding, gap filling and assembly correction with long reads"

### Supplementary Figures

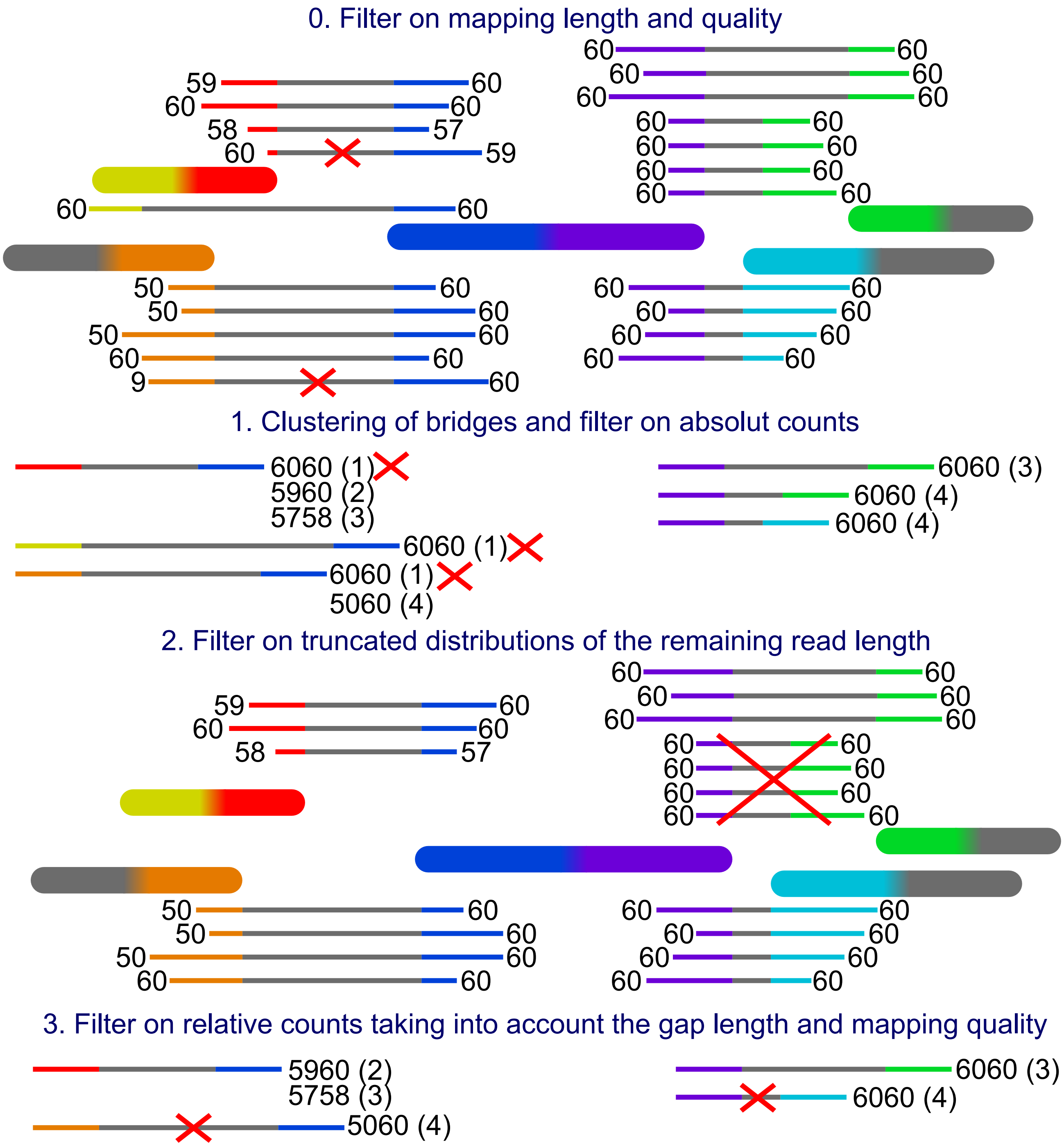


**Fig. S1 Filter example.** The big blocks with rounded corners are the contigs and the thin blocks are reads connecting them. The grey part in the middle of the reads represents the gap and the coloured parts at both ends represent the mapping to the contigs. The numbers at each end are the mapping qualities. The yellow mapping represents a mapping to the other side of the same contig as the red mapping and the green mapping has reads with two different gap lengths. The prefiltering on mapping length and quality happens on the mapping level and thus the crossed out bridges in this step will never be created. This allows the unfiltered mapping of the read to form another bridge for reads with more than two mappings. The mapping qualities are combined putting the lower quality in front and the bridges are counted cumulative from the highest to the lowest combined quality. In the second filter one bridge is removed, because the mappings are truncated on the purple side and likely stem from a different repeat copy.

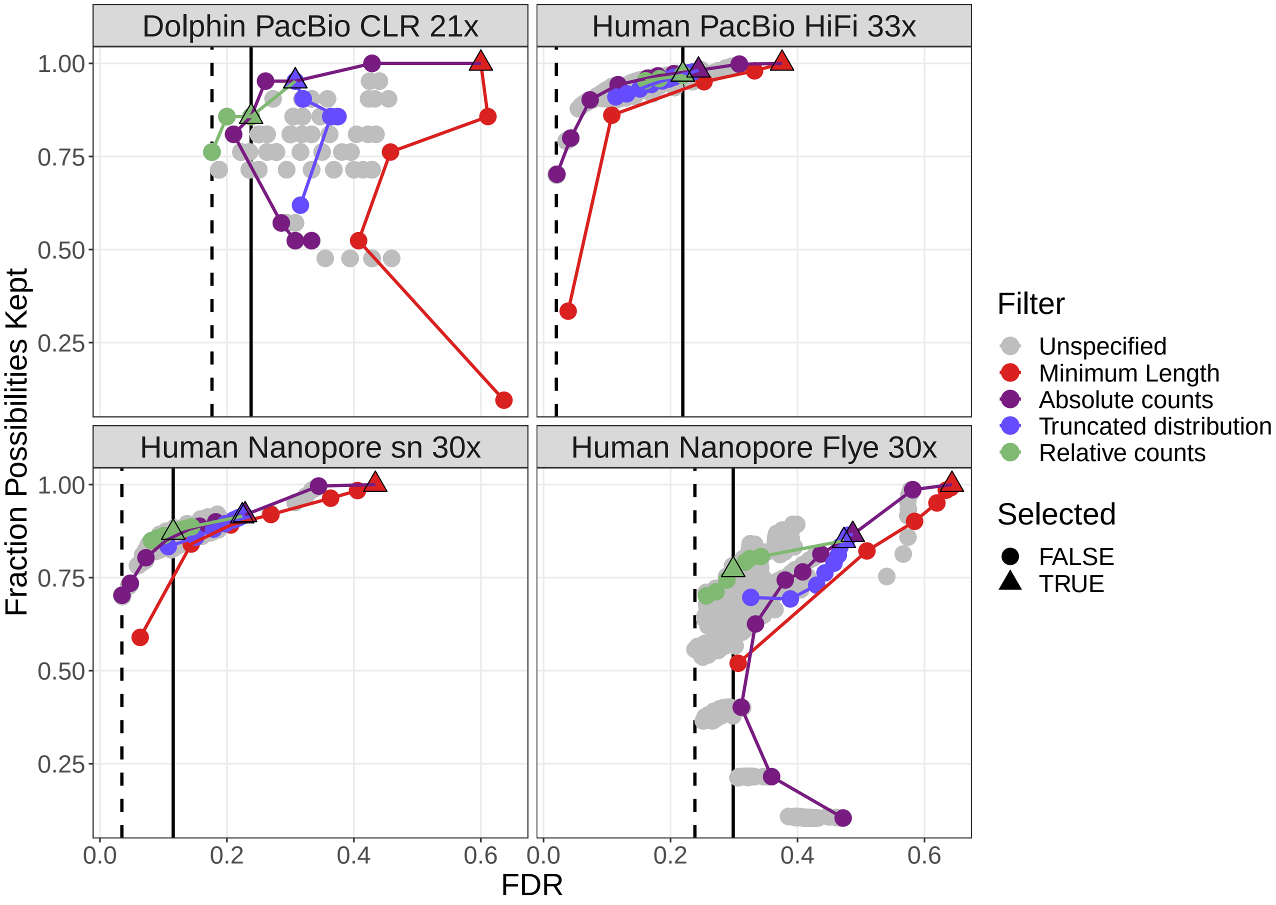


**Fig. S2 Filter performance for four datasets showing FDR and Completeness.** The filters are ordered in the legend by their order of execution. The triangles mark the selected value for a given filter, which all subsequent filters use. The unspecified category are the possible combinations of filter values tested in a grid search for which the values of the previous filters were not selected. More details can be found in figures S3 to S6. The solid vertical line marks the FDR with default parameter, the dashed vertical line marks the lowest FDR found in the grid search. The coloured lines connect the dots in order from least to most stringent for each filter.

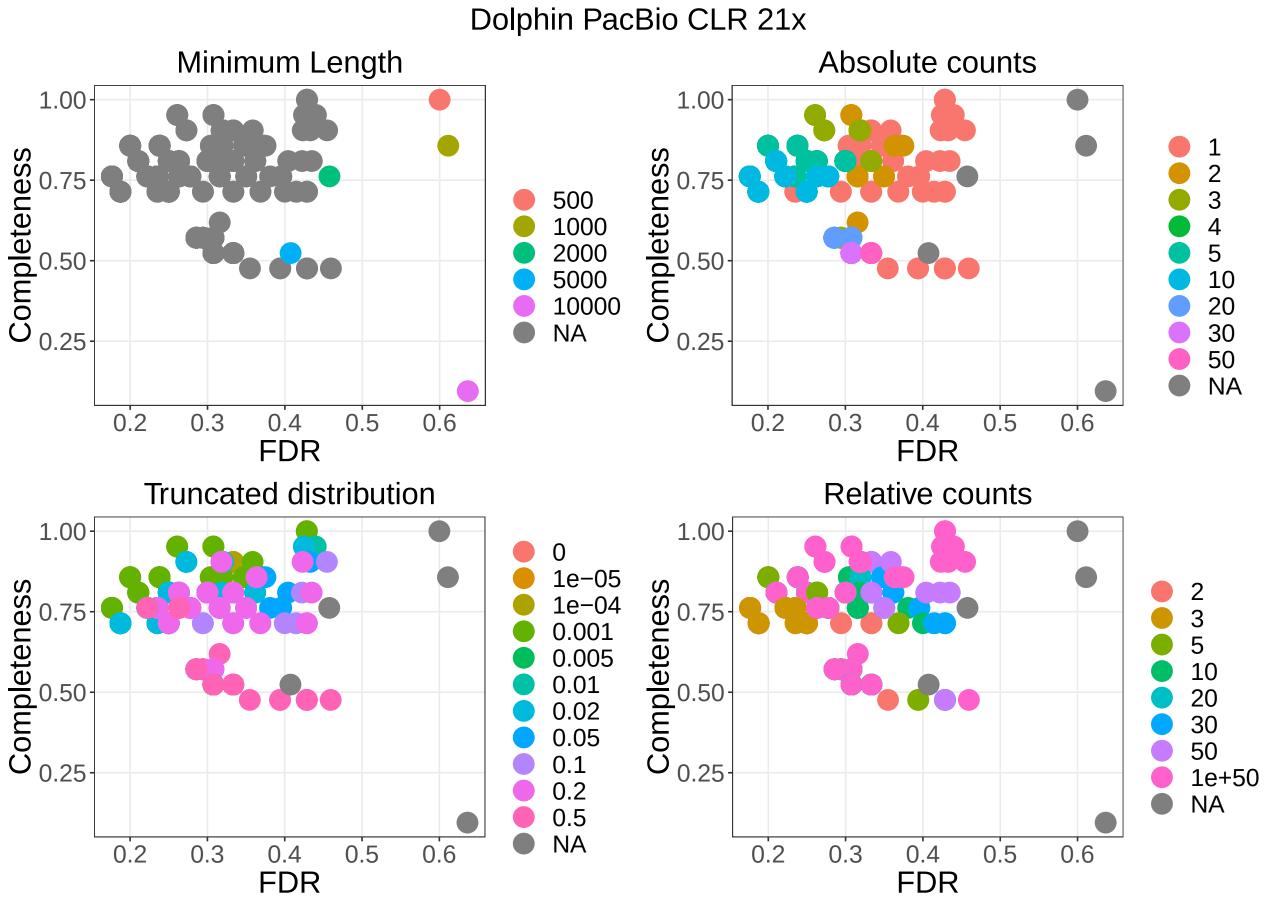


**Fig. S3** **Filter performance for the dolphin PacBio dataset with 21x coverage showing the detailed filter values for each filter.**

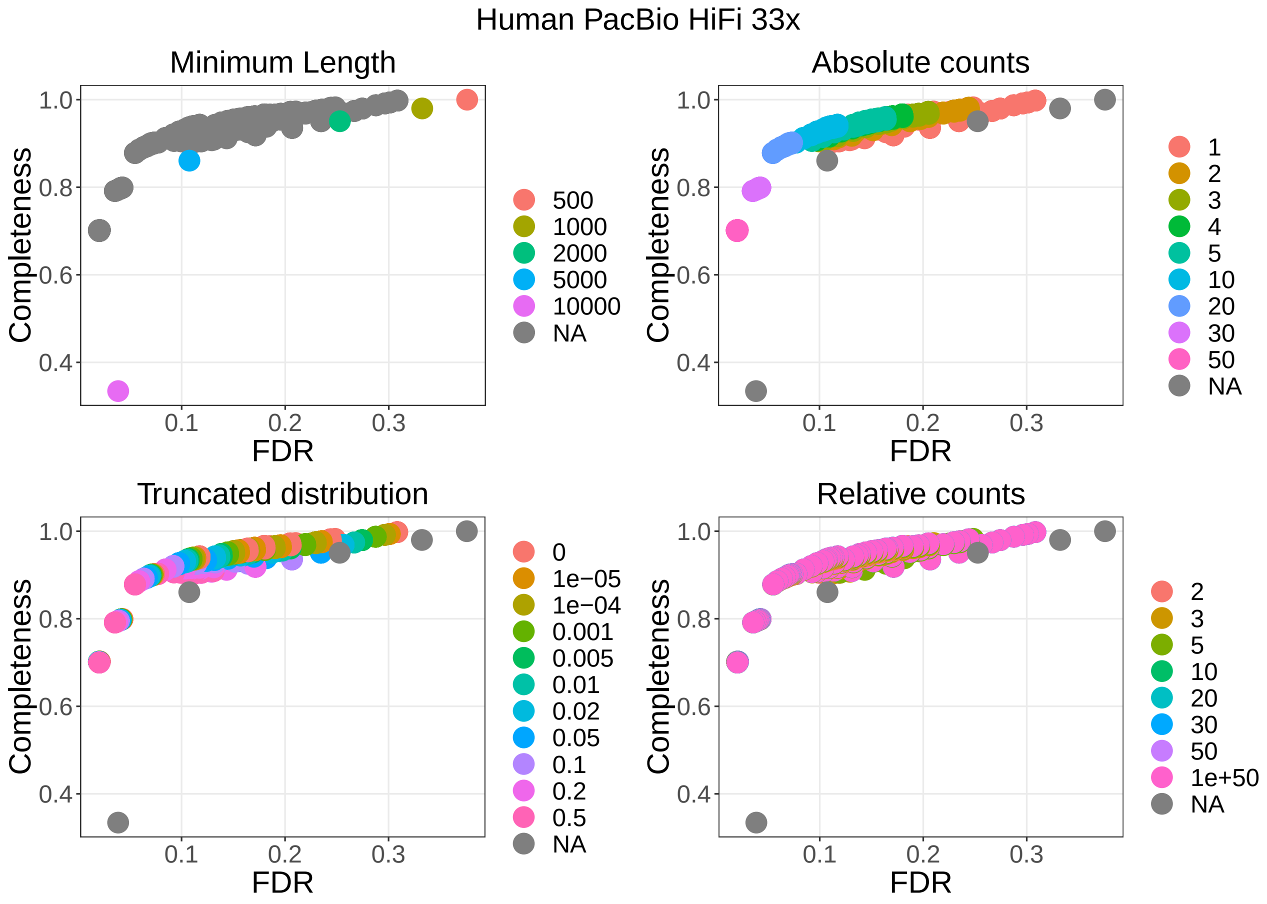


**Fig. S4** **Filter performance for the human PacBio dataset with 33x coverage showing the detailed filter values for each filter.**

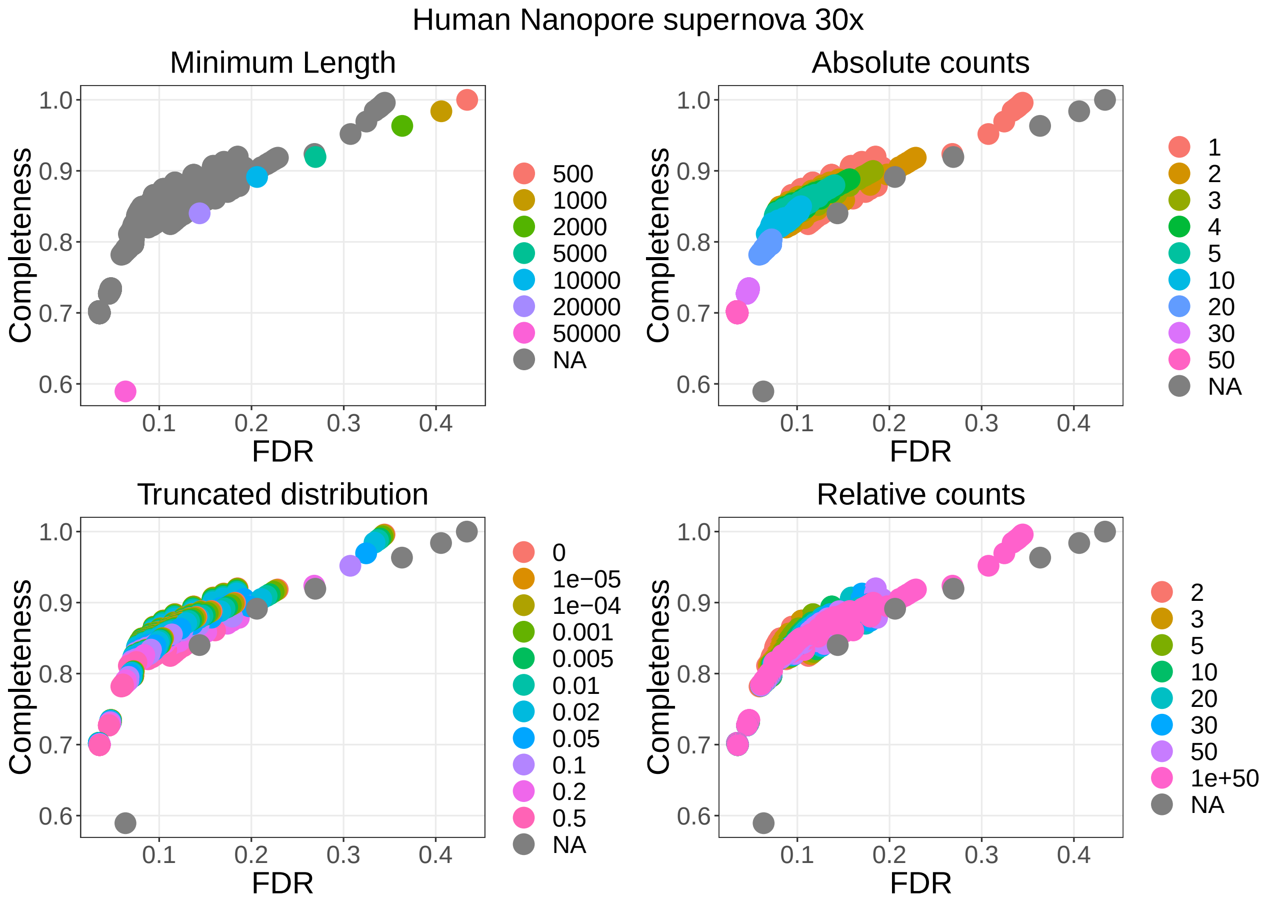


**Fig. S5 Filter performance for the human Nanopore dataset with 30 coverage starting from the supernova assembly showing the detailed filter values for each filter.**

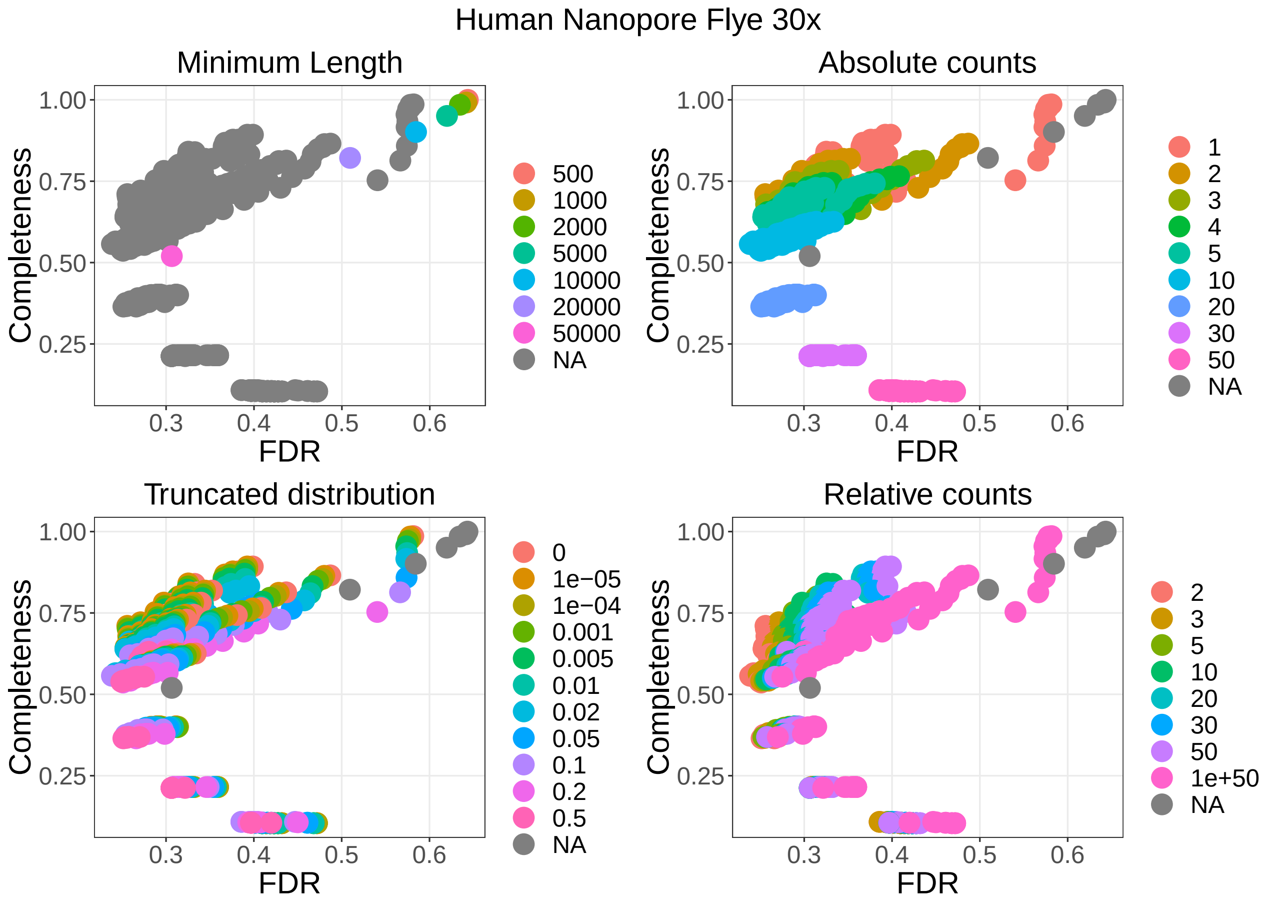


**Fig. S6 Filter performance for the human Nanopore dataset with 30x coverage starting from the Flye assembly showing the detailed filter values for each filter.**

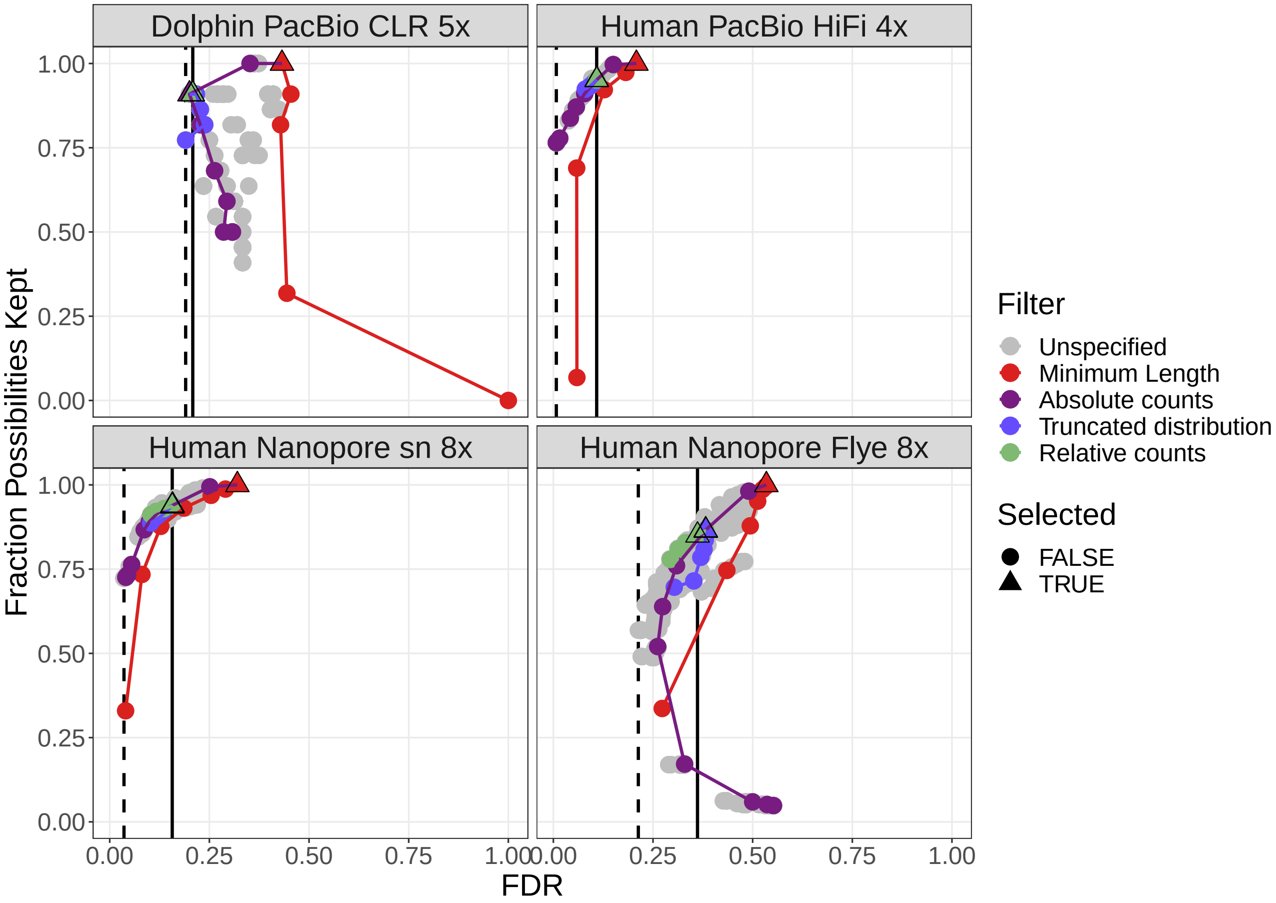


**Fig. S7 Filter performance for four datasets with very low coverage showing FDR and Completeness.** The filters are ordered in the legend by their order of execution. The triangles mark the selected value for a given filter, which all subsequent filters use. The unspecified category are the possible combinations of filter values tested in a grid search for which the values of the previous filters were not selected. More details can be found in figures S3 to S6. The solid vertical line marks the FDR with default parameter, the dashed vertical line marks the lowest FDR found in the grid search. The coloured lines connect the dots in order from least to most stringent for each filter.

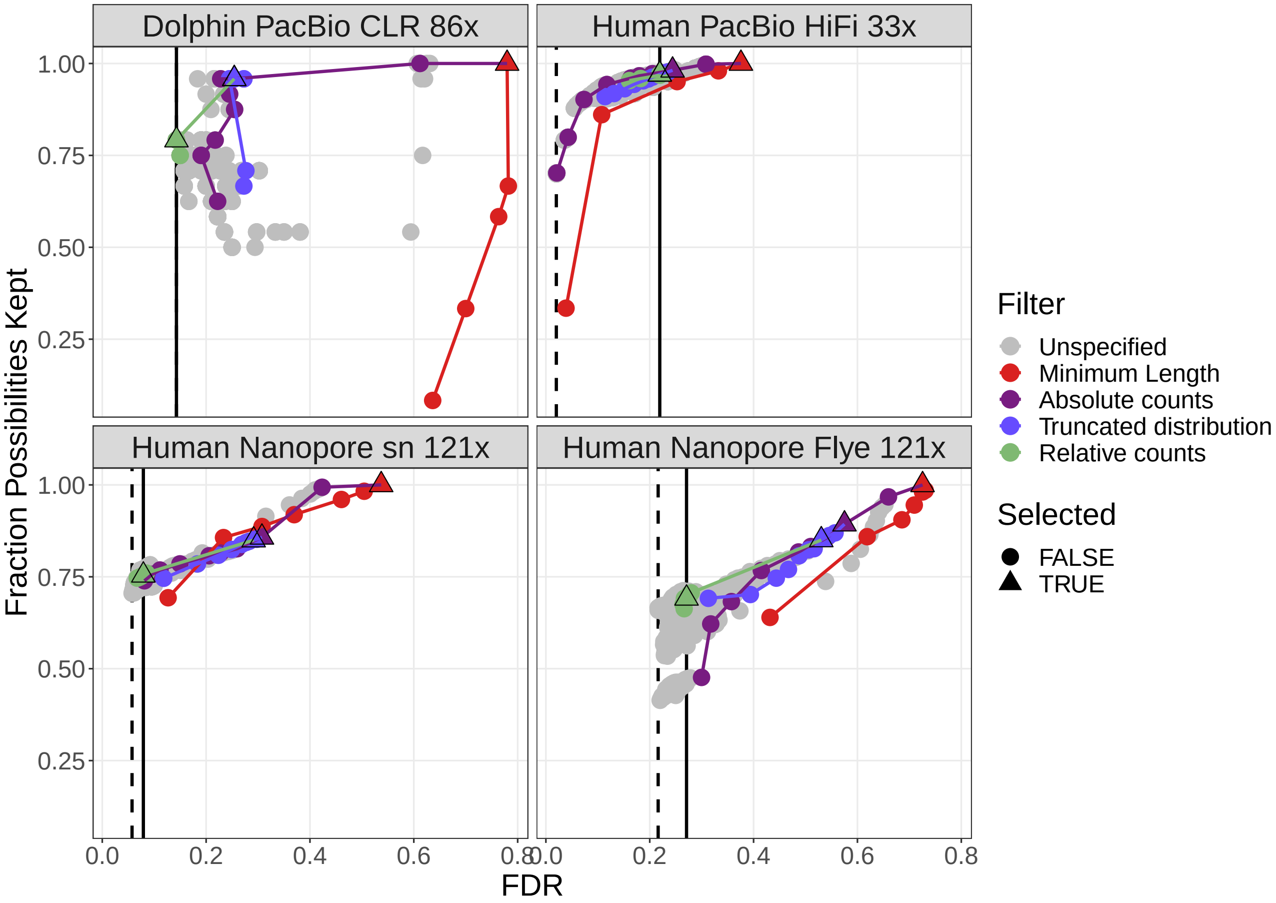


**Fig. S8 Filter performance for four datasets with very high coverage showing FDR and Completeness.** The filters are ordered in the legend by their order of execution. The triangles mark the selected value for a given filter, which all subsequent filters use. The unspecified category are the possible combinations of filter values tested in a grid search for which the values of the previous filters were not selected. More details can be found in figures S3 to S6. The solid vertical line marks the FDR with default parameter, the dashed vertical line marks the lowest FDR found in the grid search. The coloured lines connect the dots in order from least to most stringent for each filter.

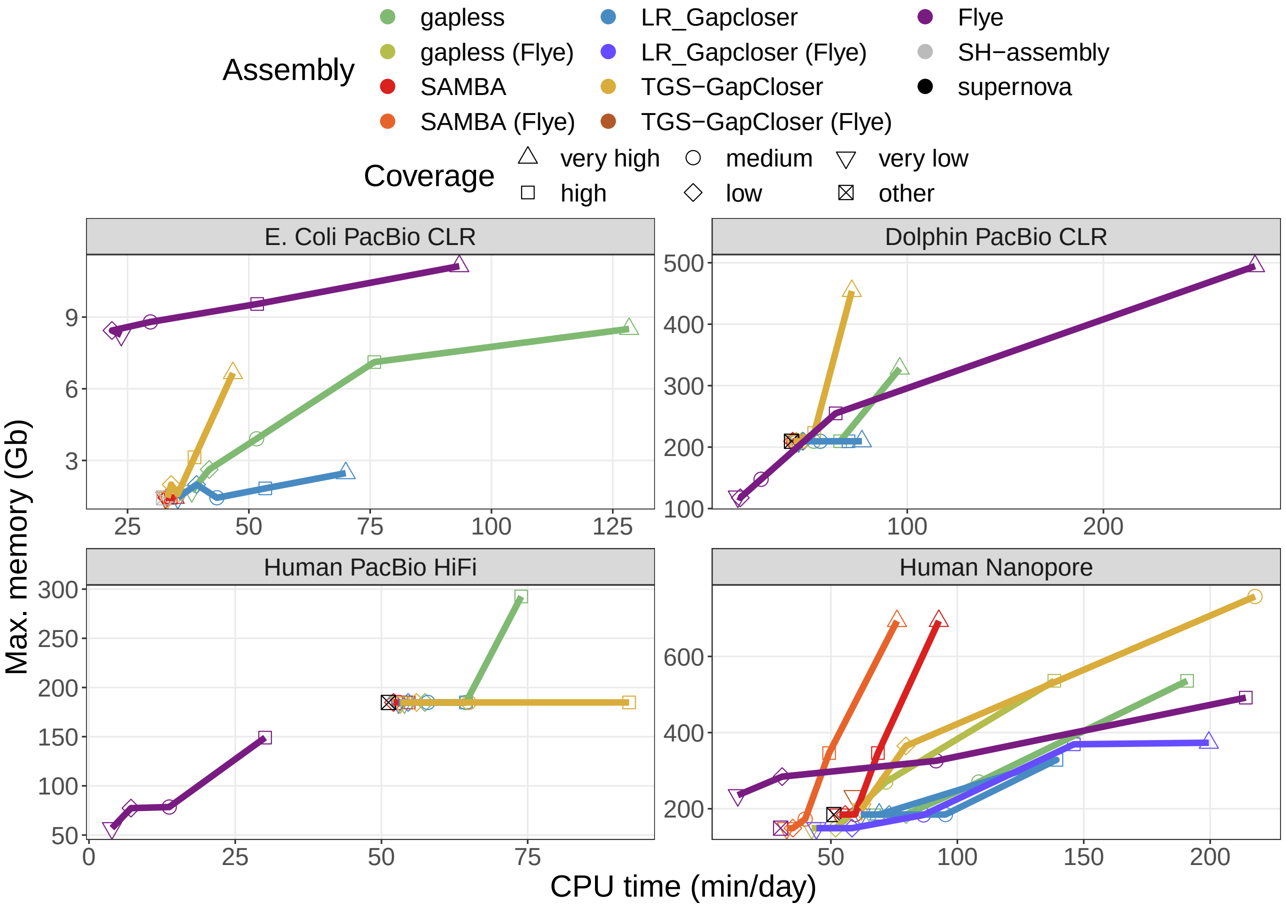


**Fig. S9 Required CPU time and memory to create the compared assemblies.** The time and memory of the base assembly were included in the values for the scaffolding results. The times for *E. coli* are specified in minutes, while the other times are given in days. The datasets are specified in Table 1 in the main manuscript. Human PacBio HiFi data do not have a very high coverage category. The "other" coverage category are the base assemblies improved with scaffolding and gap filling, which were created from a different dataset. Multiple assemblies were excluded, because they crashed or did not finish within the one week limit of the cluster. Details are given in the methods section.

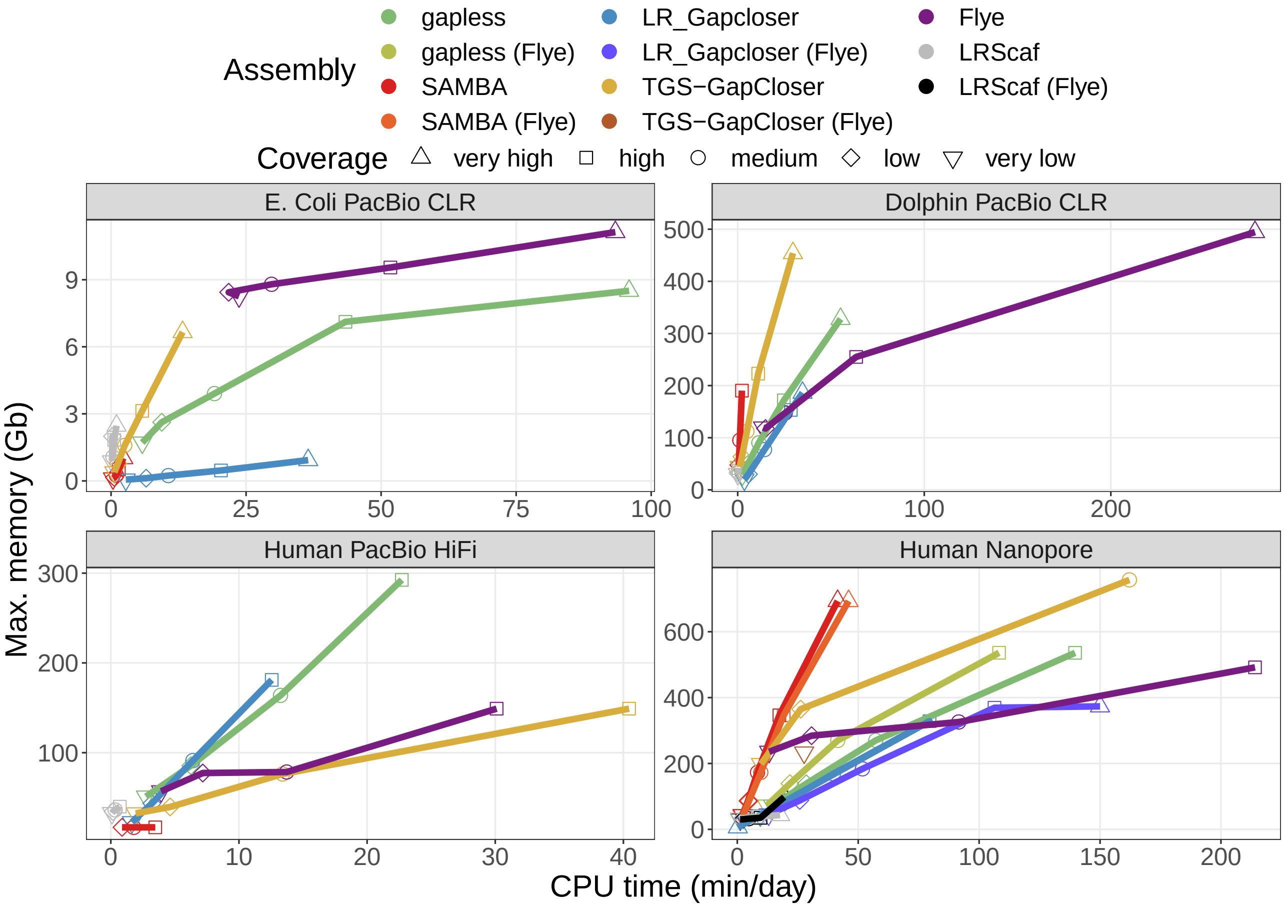


**Fig. S10 Individually required CPU time and memory for the tools used to create the compared assemblies.** The times for *E. coli* are specified in minutes, while the other times are given in days. The datasets are specified in Table 1 in the main manuscript. Human PacBio HiFi data do not have a very high coverage category. Multiple assemblies were excluded, because they crashed or did not finish within the one week limit of the cluster. Details are given in the methods section. The time and memory of the base assemblies are not included to keep colors distinguishable, but can be found in Fig. S9. LRScaf was run before LR_Gapcloser and TGS-GapCloser to perform the scaffolding.

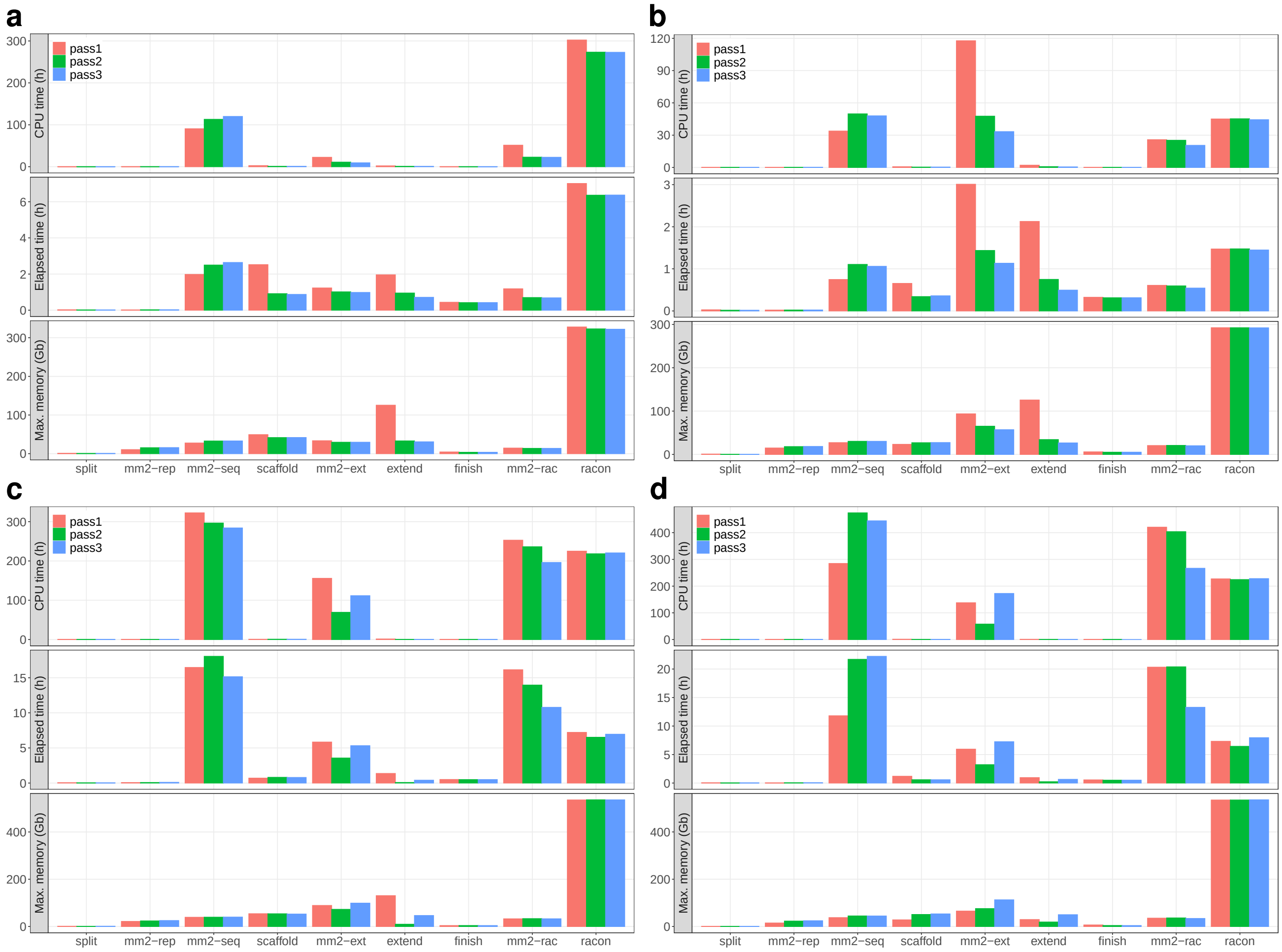


**Fig. S11 Time and memory requirements for the individual parts of the gapless pipeline.** The individual processes are **gapless** `split` (split), minimap2 aligning the assembly to itself (mm2-rep), minimap2 mapping the reads against the split assembly (mm2-seq), **gapless** `scaffold` (scaffold), minimap2 all-vs-all alignment of extending reads (mm2-ext), **gapless** `extend` (extend), **gapless** `finish` (finish), minimap2 mapping the reads against the scaffolded assembly (mm2-rac) and racon. The input data are 86x dolphin PacBio CLR (a), 33x human PacBio HiFi (b) and 61x human Nanopore starting from the **Flye** assembly (c) and **supernova** assembly (d).

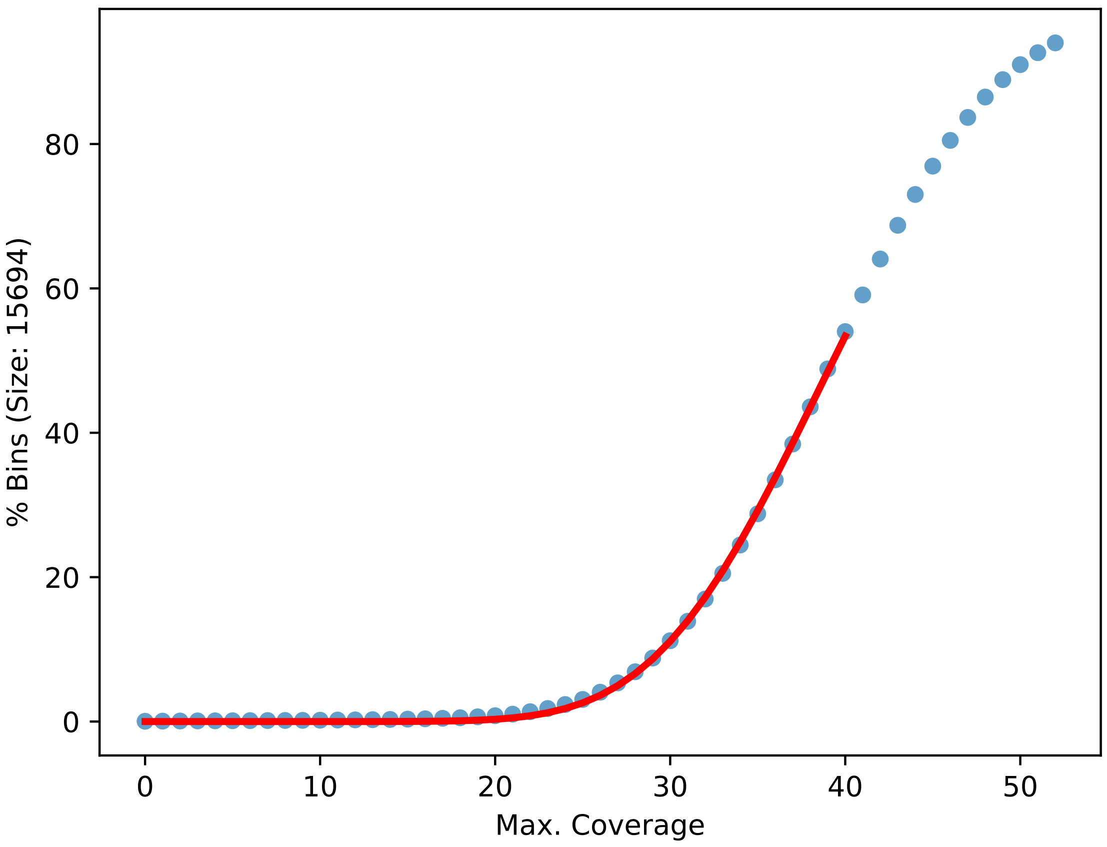


**Fig. S12 Cumulative distribution of counts over bins of size 15694 bases.** The distribution is taken from the first **gapless** iteration of the human Nanopore data with 61x coverage starting from the **Flye** assembly. The red line is a fitted negative binomial CDF.

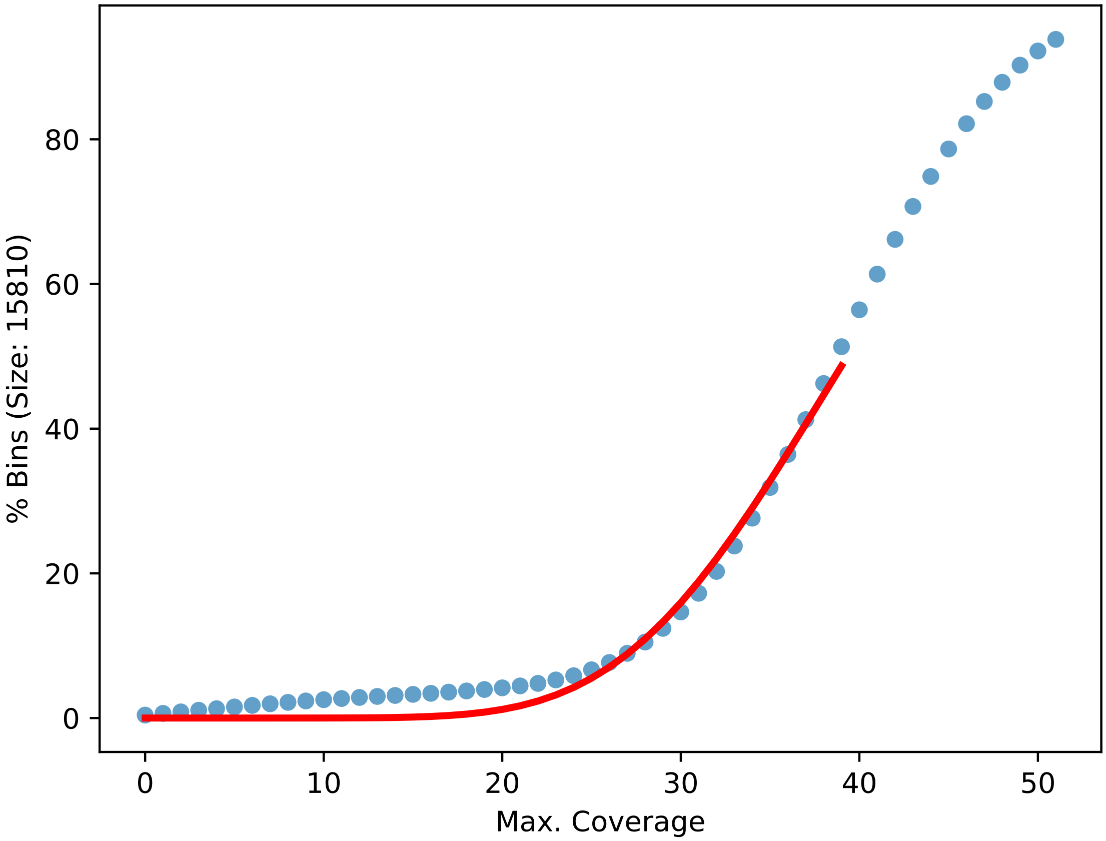


**Fig. S13 Cumulative distribution of counts over bins of size 15810 bases.** The distribution is taken from the first **gapless** iteration of the human nanopore data with 61x coverage starting from the **supernova** assembly. The red line is a fitted negative binomial CDF.

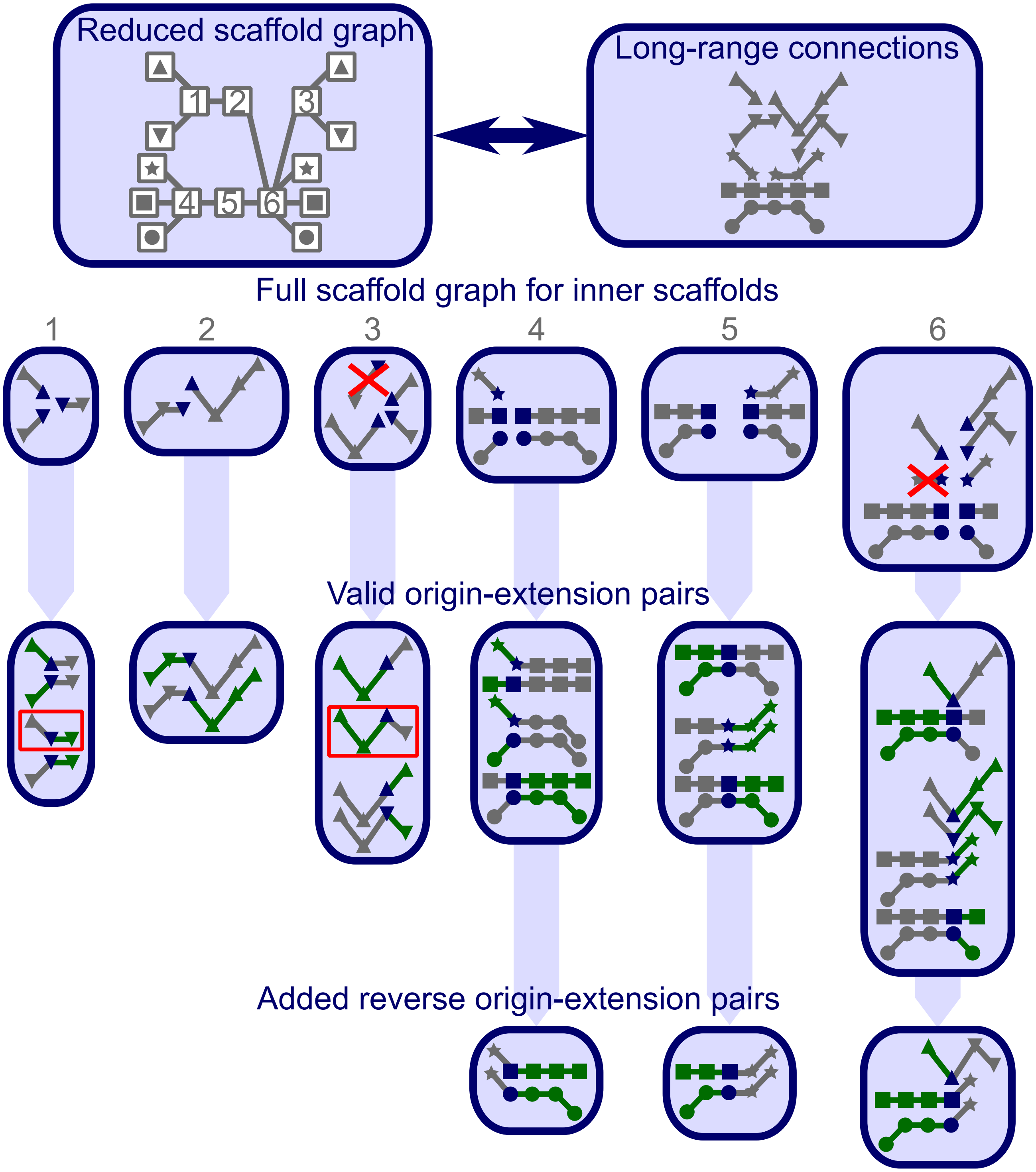


**Fig. S14 Overview on scaffold graph and origin-extension pairs.** The scaffold graph is represented in three different ways. The reduced graph shows only the connections to the next scaffold. The symbols represent the true connections between the outer scaffolds, normally unknown. Alternatively, the long-range connections are shown, marked with the symbols of the outer scaffolds they are including. To simplify the graph, no connections purely containing inner scaffolds are included in this example. The full graph representation (for the inner scaffolds labeled in the reduced graph) shows the long-range connections, split for every starting scaffold (blue). The crossed out connections are removed, because they are fully contained within another connection. The origin-extension pairs built for every starting scaffold are shown below with the origins colored in green and the extensions in grey. The pairs in red squares are only accepted, because the allowed extensions are only filtered at diverging scaffolds in the origin. In this example those scaffold sides only have one origin. Thus, they are never filtered.

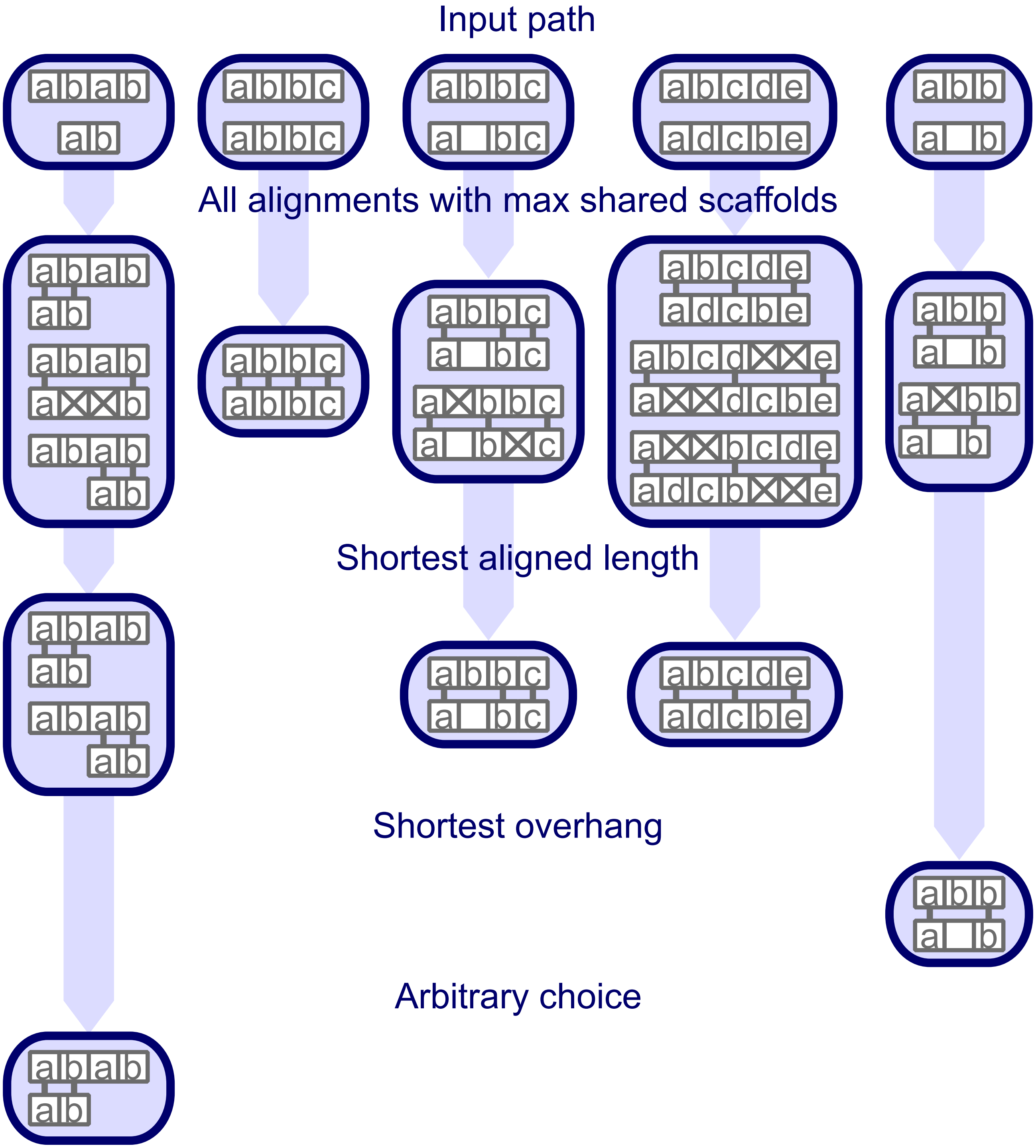


**Fig. S15 Example of pairwise path alignment.** The first row shows five examples of paths to align. Identical scaffolds are labeled with the same letter. Unlabeled scaffolds are only present in one path and cannot be aligned. The next rows show all possible alignments after a given filtering step. Aligned scaffolds are connected with a short line. Steps that do not filter alignments are not shown for the given example.
